# Two successive oligomeric Munc13 assemblies scaffold vesicle docking and SNARE assembly to support neurotransmitter release

**DOI:** 10.1101/2023.07.14.549017

**Authors:** Manindra Bera, Kirill Grushin, R Venkat Kalyana Sundaram, Ziasmin Shahanoor, Atrouli Chatterjee, Abhijith Radhakrishnan, Seong Lee, Murugesh Padmanarayana, Jeff Coleman, Frédéric Pincet, James E Rothman, Jeremy S Dittman

## Abstract

The critical presynaptic protein Munc13 serves numerous roles in the process of docking and priming synaptic vesicles. Here we investigate the functional significance of two distinct oligomers of the Munc13 core domain (Munc13C) comprising C1-C2B-MUN-C2C. Oligomer interface point mutations that specifically destabilized either the trimer or lateral hexamer assemblies of Munc13C disrupted vesicle docking, trans-SNARE formation, and Ca^2+^-triggered vesicle fusion in vitro and impaired neurotransmitter secretion and motor nervous system function in vivo. We suggest that a progression of oligomeric Munc13 complexes couples vesicle docking and assembly of a precise number of SNARE molecules to support rapid and high-fidelity vesicle priming.

## Introduction

Chemical synaptic transmission is an orchestrated sequence of molecular events that endows synapses with a remarkably fast and tunable communication channel through precise regulation of synaptic vesicle (SV) fusion. The low basal rate of SV fusion at a release site in the absence of a Ca^2+^ trigger can jump by a factor of ten to one hundred million upon local Ca^2+^ elevation, and the latency between Ca^2+^ arrival and neurotransmitter release can be shorter than 100 µsec ^1–3^. This extraordinary performance distinguishes SV exocytosis from other cellular trafficking events and requires a specialized machinery. At its core, this process is driven by a tightly regulated assembly of SNARE proteins along with several critical SNARE-associated proteins including Munc13-1, Munc18-1, Synaptotagmin 1 (Syt1), and Complexin among others ^4^. At the mechanistic level, many questions remain largely unanswered: How many SNAREs typically assemble to drive SV fusion and how is this number supervised? How is a vesicle brought into proximity with the plasma membrane while full SNARE assembly and vesicle fusion are prevented in the absence of a calcium trigger? How do environmental signals such as residual Ca^2+^ and lipids influence the steps leading to SV priming and fusion? Residing at the hub of the presynaptic release apparatus, Munc13-1 is involved in nearly every aspect of presynaptic function including release site building, synaptic vesicle (SV) docking and priming, SNARE assembly, SV fusion, and use-dependent plasticity ^4,5^. While much has been learned about Munc13-1 structure and function since its discovery over 30 years ago, these functions of Munc13-1 still lack a thorough molecular understanding. Moreover, many of the questions raised above are directly related to Munc13-1 function. We previously uncovered two novel oligomeric assemblies of Munc13-1 and proposed a model based on their structures that provides insight into these questions and the distinct roles of Munc13-1 in promoting SV capture and the steps leading to vesicle priming ^6^.

Distinct homo-oligomeric assemblies of the Munc13-1 core domain (C1-C2B-MUN-C2C, hereon referred to as Munc13C) were observed in vitro via cryo-electron tomography (Cryo-ET) of two-dimensional protein crystals self-assembling between two phospholipid bilayers. These were proposed to correspond to sequential states of SV docking and pre-priming so that SVs are maintained at precise distances from target membranes while SNARE assembly is gated by the coordinated occlusion of SNARE-binding sites on the MUN domain ^6^. In this scenario, the C-terminal end of Munc13C assists in SV capture and tethering while the N-terminal end interacts with the plasma membrane and responds to elevated DAG and intracellular Ca^2+^ to transition the upright trimer state (hexamer of trimers) to the lateral (hexamer of monomers) state.

Several activities could be accomplished by this proposed Munc13C topology and sequence of events. We suggested that SVs are captured and tethered in the upright state only when a critical mass of Munc13C monomers have pre-assembled ^7^. The upright state can then transition to the lateral state bringing the SV nearer to the plasma membrane while also preventing MUN-catalyzed SNARE assembly due to occlusion of the MUN domain SNARE binding sites by neighboring monomers. This transition is proposed to be promoted by a translocation of the C1 domain in the presence of diacylglycerol (DAG). Finally, ratcheting of the MUN domains down onto the plasma membrane (potentially assisted by Ca^2+^ ions binding the C2B domain) would catalyze the simultaneous assembly of precisely 6 SNAREpins (based on the six-fold symmetry of the lateral hexamer) while helping to bring the SV and plasma membranes into close proximity for SV priming. This model specifically addresses aspects of the questions posed above.

In the present study, we designed and investigated a small number of predicted interface-disrupting point mutations and used a variety of structural, biochemical, and in vivo approaches to test this model. Disruptions in either a MUN-MUN interface in the trimer or in a MUN-C2C interface in the lateral hexamer led to pronounced structural alterations of Munc13C as well as severe defects in vesicle docking, MUN-catalyzed SNARE assembly, and Ca^2+^-triggered fusion in vitro as well as impaired neurotransmitter secretion in vivo. Specifically, wild-type Munc13C promoted the rapid assembly of precisely six SNAREpins mirroring the six-fold symmetry of the lateral hexamer, whereas Munc13C structural variants could trigger at most one or two stable SNAREpins to form. These observations highlight the importance of two oligomeric Munc13 assemblies in coupling vesicle docking to a fixed number of assembled SNAREpins and provide structural and functional insights into the kinetics and fidelity of vesicle priming.

## Results

### Disruption of the Munc13C trimer interface

We recently reported two markedly distinct oligomeric states of Munc13C designated ‘upright trimers’ and ‘lateral hexamers’ based on their respective arrangements and symmetries in crystals formed between lipid bilayers as determined by cryo-electron tomography (CryoET) ^6^. In the upright trimer arrangement, 6 trimers of Munc13C tethered two lipid bilayers with a spacing of 21 nm, consistent with a synaptic vesicle (SV) held in a pre-docked state with no SNARE assembly (**Fig. 1A**). Due to the low resolution of the 3D reconstruction in the trimer interface region, the specific amino acids that form this interface could not be identified with high confidence. However, the surface charge distribution suggested that the interface is electrostatic in nature, possibly corresponding to residues D1358 and D1369 in helix H12 of MUN-D subdomain and residues K1494, K1495, and K1500 in helix H15 of MUN-D subdomain (**Fig.1A & B**) using the helix nomenclature of Rizo and colleagues and rat Munc13-1 amino acid numbering ^8^. Protein sequence alignments for H12 and H15 across metazoa revealed conservation of the acidic residues in H12 as well as the lysines in H15 (**Fig. 1C**).

**Figure 1.**
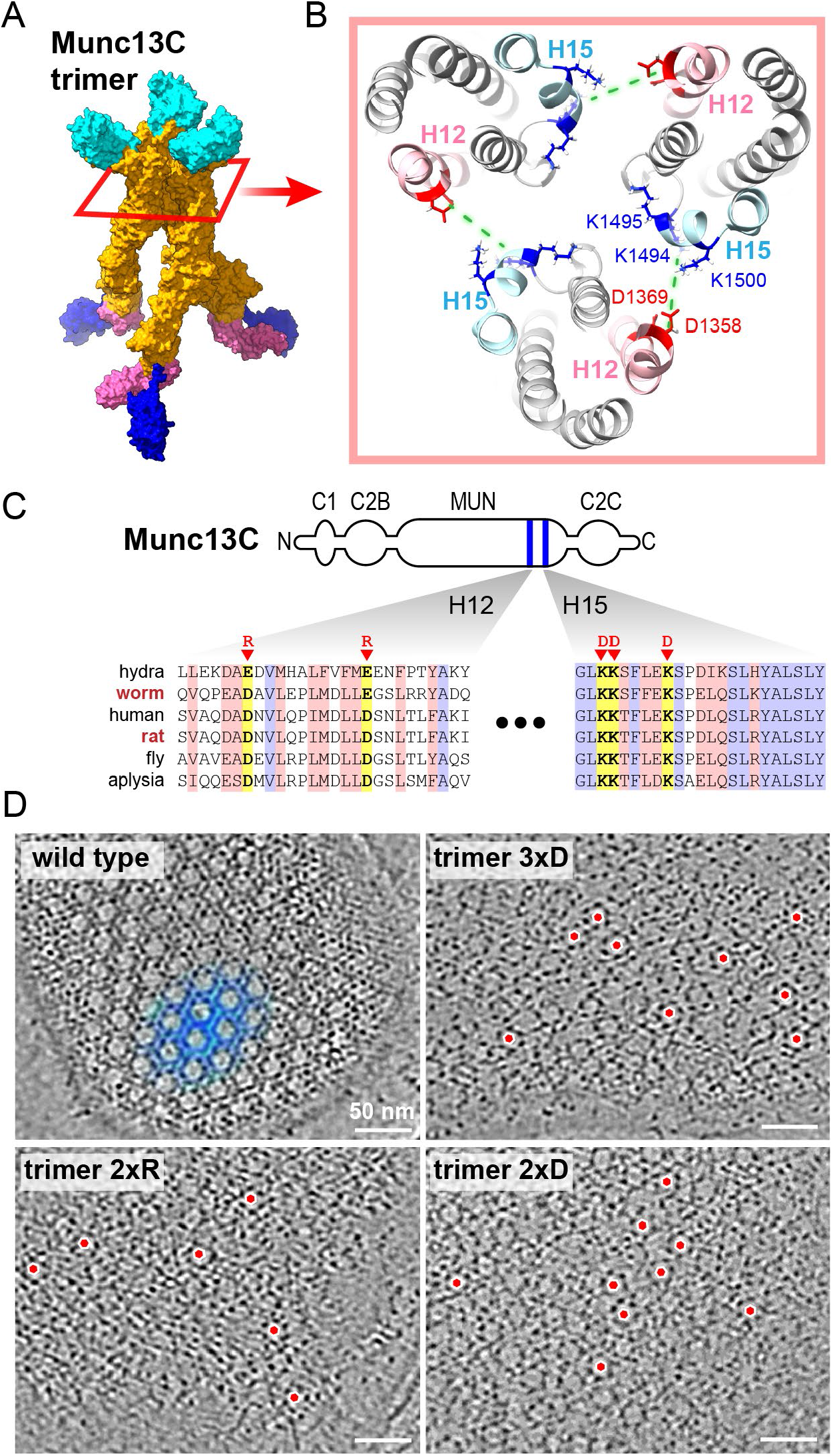
Disruption of the Munc13C trimer interface. **A.** Surface representation of the Munc13C trimer (PDB: 7T7R). Domain color code: C1 (*pink*), C2B (*blue*), MUN (*orange*), and C2C (*cyan*). **B.** Top view of the trimer interface. Green dashed lines show the proposed interaction between acidic (*red*) and alkaline (*blue*) residues within the MUN-D region of three Munc13C molecules. Putative residues involved in trimer formation are D1358 and D1369 (helix H12 of MUN domain) and K1495 and K1500 (helix H15 of MUN domain). **C.** Sequence alignments for MUN H12 and H15 helices for a select group of metazoa with identical and similar residues highlighted in blue and red, respectively. The residues mutated in the trimer variants are highlighted in yellow. **D.** Top view of representative sections from cryo-electron tomograms showing effect of mutations on crystal formation by Munc13C between lipid bilayers. Wild type Munc13C (*top left*) forms a continuous hexagonal lattice (*blue*) whereas reversing polarity of K1494/K1495/K1500 (3xD *top right*), D1358/D1369 (2xR *lower left*) and K1495/K1500 (2xD *lower right*) disrupted the lattice. Isolated hexagons (*red circles*) are visible without forming the characteristic honeycomb pattern of wild-type Munc13C. Scale bars are 50 nm. Protein sequences used for alignment: hydra = *H. vulgaris* (XP012567200), worm = *C. elegans* (NP001021874), human = *H. sapiens* (NP001073890), rat = *R. rattus* (XP032775345), fly = *D. melanogaster* (NP651949), aplysia = *A. californica* (XP035824675).

To test the functional significance of the upright trimer interface, two Munc13C variants harboring the following charge reversals were designed to produce electrostatic repulsion within the interface: D1358R/D1369R (trimer 2xR) and K1494D/K1495D/K1500D (trimer 3xD). Notably, these protein variants were stable and well-folded. The interface variants were combined with negatively charged phospholipid membranes under the same conditions that generate hexagonal crystals with wild-type Munc13C (**Fig. 1D**) and assessed for gross structural alterations by CryoET. Notably, both the trimer 2xR and 3xD mutants disrupted continuous hexagonal lattice formation apparent in the CryoET top view (**Fig. 1C**). Isolated lateral hexagons were observed between two lipid bilayers along with scattered unorganized Munc13C monomers in the interface variants. Hexagonal lattices were rarely observed and only occurred in small patches, indicating that the lattice organization enabled by the upright trimers was largely abolished. To better determine which basic residues were required for the trimer interface, two more Munc13C variants were created with only a pair of lysine charge reversals: K1495D/K1500D and K1494D/K1495D. Whereas the K1494D/K1495D variant did not alter the hexagonal crystal lattice, the K1495D/K1500D variant (hereon referred to as trimer 2xD) was similar to the 3xD variant, consistent with K1495 and K1500 participation at the trimer interface. The tomograms of all four Munc13C timer interface variants appeared similar in side-view cross section with a pronounced linear protein density centered between bilayers, indicative of an intact lateral hexamer assembly (**Fig. S1A-D**). Moreover, the bilayer separation within those regions was identical to wild type (21 nm), indicating the presence of a vertical conformation of Munc13C monomers. Hence, charge reversals in H12 and H15 of the MUN domain selectively destabilized the upright trimer assembly.

### Disruption of the Munc13C lateral interface

The other Munc13C oligomeric arrangement observed by CryoET was a lateral hexamer corresponding to an intermembrane spacing of about 14 nm, and the protein interface within this structure was formed by the C2C domain of one monomer and the MUN domain of its nearest neighbor (**Fig. 2A**) ^6^. Previously we posited two interface regions from this structure. First, a hydrophobic region was observed between C2C and MUN-C (F1176 and I1215 on H7 and H8 respectively, and W1626 and W1684 in C2C, **Fig. 2B**). Second, a polar region was observed between MUN-B and C2C (D1039 near H3 and E1139 in H6 of the MUN domain and R1673 and R1678 on the C2C domain, **Fig. 2B**). Sequence alignments across metazoa indicated strong conservation of the nonpolar interface (**Fig. 2C**), whereas the electrostatic interface displayed higher variability across phylogeny (**Fig. S1B**).

**Figure 2.**
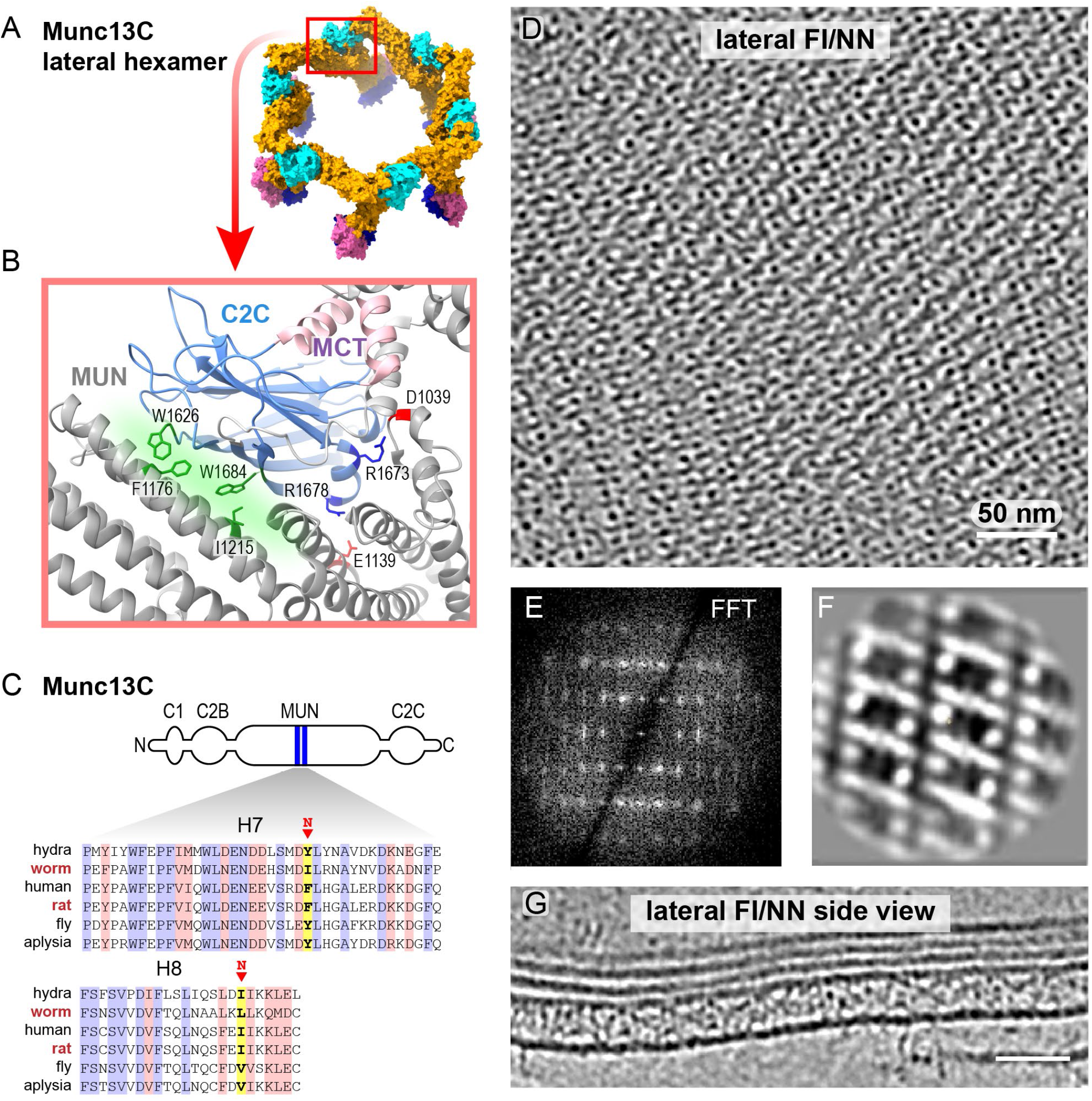
Disruption of the Munc13C lateral interface. **A.** Surface representation of the Munc13C lateral hexamer (PDB: 7T7C). Domain color code: C1 (*pink*), C2B (*blue*), MUN (*orange*), C2C (*cyan*). **B.** Side view of the lateral interface. C2C (*blue*) and MUN-C terminal helices (MCT, *light pink*) are positioned above the area of MUN-B and MUN-C domains (gray) of a neighboring Munc13C monomer. In detail, C2C contacts H7, H8 of MUN-C as well as H4, H6, and the loop between H3 and H4 of MUN-B (*green highlight*). Proposed residues involved in the lateral interface are labeled. **C.** Sequence alignments for MUN H7 and H8 helices with identical and similar residues highlighted in blue and red, respectively. The residues mutated in the lateral FI/NN variant are highlighted in yellow. Protein sequence accession numbers are given in Figure 1. **D.** Representative section of a cryo-electron tomogram with a top view of crystals formed by Munc13C F1176N/I1215N mutant between lipid bilayers. **E.** Fast Fourier Transform (FFT) of the cryo-tomogram indicates a 2-fold rectangular symmetry for this crystal. **F.** A slice through a 3D volume of the initial low-resolution reconstruction. **G.** Section through a cryo-electron tomogram with a side view of the lateral FI/NN crystal. Note the absence of a characteristic density between the bilayers (**Figure S1A**), indicating that the lateral hexamer is not present. Distance between bilayers is similar to wild type (∼21 nm). Scale bars are 50 nm.

Based on the higher conservation of the predicted nonpolar interface, two bulky residues (F1176 and I1215 in H7 and H8 respectively) were mutated to aspargines (hereon referred to as the lateral FI/NN variant) and this variant was examined for disruptions in the hexagonal lattice via CryoET. For some in vitro experiments, we also generated a quadruple substitution including D1039R/E1139R (hereon referred to as the lateral quad variant). The tryptophan residues on C2C domain were not substituted because these changes may impair C2C-membrane binding and thus and indirectly impair crystal formation. Interestingly, the lateral FI/NN variant disrupted the Munc13C hexagonal lattice while also generating a novel rectangular crystal lattice (**Fig. 2C**). Qualitatively, the Munc13C density was higher than wild type with visible rows of spots and lines probably corresponding to vertical and lateral orientation of Munc13 molecules, respectively. The lateral hexagon oligomer was likely eliminated since there were no detectable hexagonal lattices in the CryoET top view (**Fig. 2D**), and lateral FI/NN crystals lacked the characteristic linear density in the side view (**Fig. 2G** compared to **Fig. S1A**). The inter-bilayer distance was 21 nm, similar to wild-type Munc13C, suggesting the presence of upright protomers in the new crystal. A preliminary low-resolution 3D reconstruction (**Fig 2E-F**) reflected some features of the crystallographic pattern consistent with vertical assemblies of Munc13C monomers with 2-fold symmetry, but its resolution was not sufficient to permit a detailed Munc13C structural modeling. In summary, the lateral FI/NN variant and the two trimer variants selectively disrupted the assembly of lateral hexamers and upright trimers, respectively.

### The trimer and lateral interface variants impair vesicle capture and fusion in vitro

Several in vitro approaches have demonstrated roles for Munc13C in vesicle tethering and fusion ^7,9–12^. We recently developed a reconstituted single-vesicle fusion assay that recapitulates both docking and priming aspects of Munc13 function in the context of several key synaptic proteins involved in SNARE assembly and fusion including Munc18-1/Syntaxin 1A, Synaptotagmin 1 (Syt1), and Complexin 1 (Cpx1) as illustrated in **Fig. 3A** ^13–15^. Briefly, the pre-assembled Munc18-1/Syntaxin 1A heterodimer and palmitoylated SNAP25 were reconstituted into fluorophore-labeled lipids (using ATTO 465-DOPE) used to form giant unilamellar vesicles (GUVs), which were burst on silicon grids to form suspended lipid bilayers along with the soluble diacylglycerol mimetic 1,2-hexanoyl-sn-glycerol (DHG) as described previously ^13^. Fluorophore-labeled vesicles (ATTO647N-DOPE) containing VAMP2 and Syt1 were then added along with Cpx1 and Munc13C in the absence of Ca^2+^ while monitoring fluorescence at the suspended bilayer via confocal microscopy.

**Figure 3.**
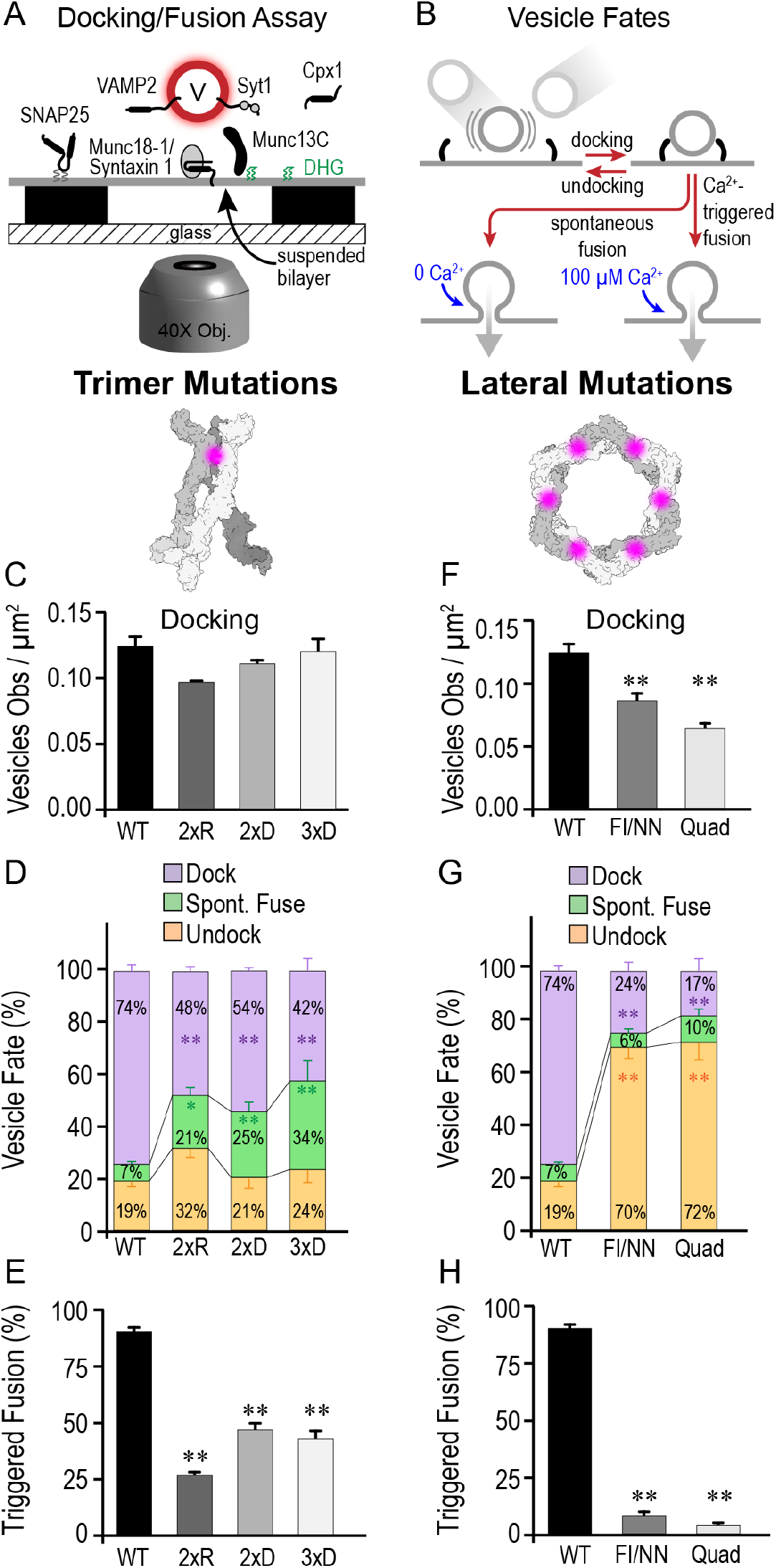
The trimer and lateral interface variants impair vesicle capture and fusion in vitro. **A.** Schematic of the single vesicle docking/fusion assay using confocal microscopy. To reconstitute Munc13/Munc18-dependent vesicle priming and fusion, palmitoylated SNAP25, 1:1 Munc18-1/Syntaxin 1 complexin, and 1,2-hexanoyl-sn-glycerol (DHG) were incorporated into the suspended lipid bilayer (labeled with 1% ATTO465-DOPE). VAMP2 and Synaptotagmin 1 (Syt1) were incorporated into small unilamellar vesicles (V) and labeled with 2% Atto647N-DOPE to visualize vesicles near the suspended bilayer. Munc13C and complexin 1 (Cpx1) were added in solution. **B.** Single vesicles were tracked by confocal microscopy and classified as docked or undocked based on their lateral mobility. Docked vesicles sometimes fused in the absence of Ca^2+^ (spontaneous fusion), whereas most vesicles fused upon addition of 100 µM Ca^2+^ (Ca^2+^-triggered fusion). Munc13C variants harboring trimer interface mutations (trimer 2xR, 2xD, and 3xD) were compared to wild type Munc13C (WT) by quantifying the number of docked vesicles (**C**), vesicle fate in the absence of Ca^2+^ (**D**), and response to 100 µM Ca^2+^ (**E**). Lateral hexamer Munc13C variants (lateral FI/NN and quad) were compared to wild-type Munc13 in **F-H**. After initial docking, vesicle fates were subdivided into three categories: vesicles remained docked (*purple*), spontaneously fused (*green*), or undocked (*orange*) (**D/G**). All data presented as mean ± standard deviation from 3 to 5 experiments. Data were compared using one-way ANOVA and the Tukey-Kramer analysis for multiple comparisons. ** *p < 0.01,* * *p < 0.05* compared to wild type.

Using this approach with the trimer and lateral interface variants described above, four aspects of Munc13C were investigated: vesicle docking, vesicle undocking, spontaneous fusion in the absence of calcium, and calcium-triggered vesicle fusion (**Fig. 3B**). Labeled vesicles diffused to the suspended bilayer and became tightly membrane-associated with minimal lateral mobility (hereon referred to as docking). The Munc13C trimer variants only weakly impacted vesicle docking (**Fig. 3C**), and docked vesicles eventually undocked with similar probabilities as observed with wild-type Munc13C (**Fig. 3D**). Surprisingly, spontaneous fusion in the absence of Ca^2+^ was significantly elevated by more than 3-fold in all trimer variants (**Fig. 3D**). Moreover, Ca^2+^-triggered fusion was impaired by about 2-fold in the 2xD and 3xD variants and by more than 3-fold in the 2xR variant (**Fig. 3E**). Thus, destabilizing the trimer interface both enhanced spontaneous fusion and impaired Ca^2+^-triggered fusion. These observations were compared with the lateral FI/NN and quad variants. In contrast to the trimer mutations, both lateral mutations significantly disrupted vesicle docking while exhibiting a 4-fold enhancement in undocking probability (**Fig. 3F&G**). And finally, while spontaneous fusion was unaffected by the lateral interface mutations, Ca^2+^-triggered fusion of docked vesicles was almost completely abolished (**Fig. 3G&H**).

### Six stable SNAREpins are assembled under clamped vesicles in the presence of Munc13C

Based on the observations described above, several questions remained unanswered. In particular, we wondered if assembled trans-SNARE complexes (hereon referred to as SNAREpins) were formed on docked vesicles. And if so, are Munc13C oligomers prerequisite for SNAREpin formation or initial vesicle capture or both? Does the underlying six-fold symmetry of the Munc13C oligomer constrain the number of SNAREpins assembled? We previously hypothesized that the lateral hexamer ratchets down onto the plasma membrane allowing exposed MUN domains to catalyze formation of central SNAREpins while bringing the membranes into closer proximity ^6^. Destabilizing early steps in a sequential process could impair downstream SNAREpin assembly, thereby accounting for the poor performance of the trimer and lateral variants in promoting Ca^2+^-triggered fusion ^16^.

To answer these questions, we recently developed a TIRF microscopy approach to monitor vesicle docking onto suspended membranes on a silicon surface with single-molecule precision (**Fig. 4A-B**) (Bera 2023). To detect SNAREpin formation on docked vesicles, we exploited the fact that Cpx1 tightly binds with the SNARE complex in a 1:1 molar ratio through its central helix ^17,18^. Specifically, Cpx1 binds within a groove formed by VAMP2 and Syntaxin 1A after the SNARE monomers have zippered partially together, and therefore detection of a persistent fluorophore signal from a single labeled Cpx1 molecule at a docked vesicle within the TIRF field could be used as a proxy for a single stable SNAREpin.

**Figure 4.**
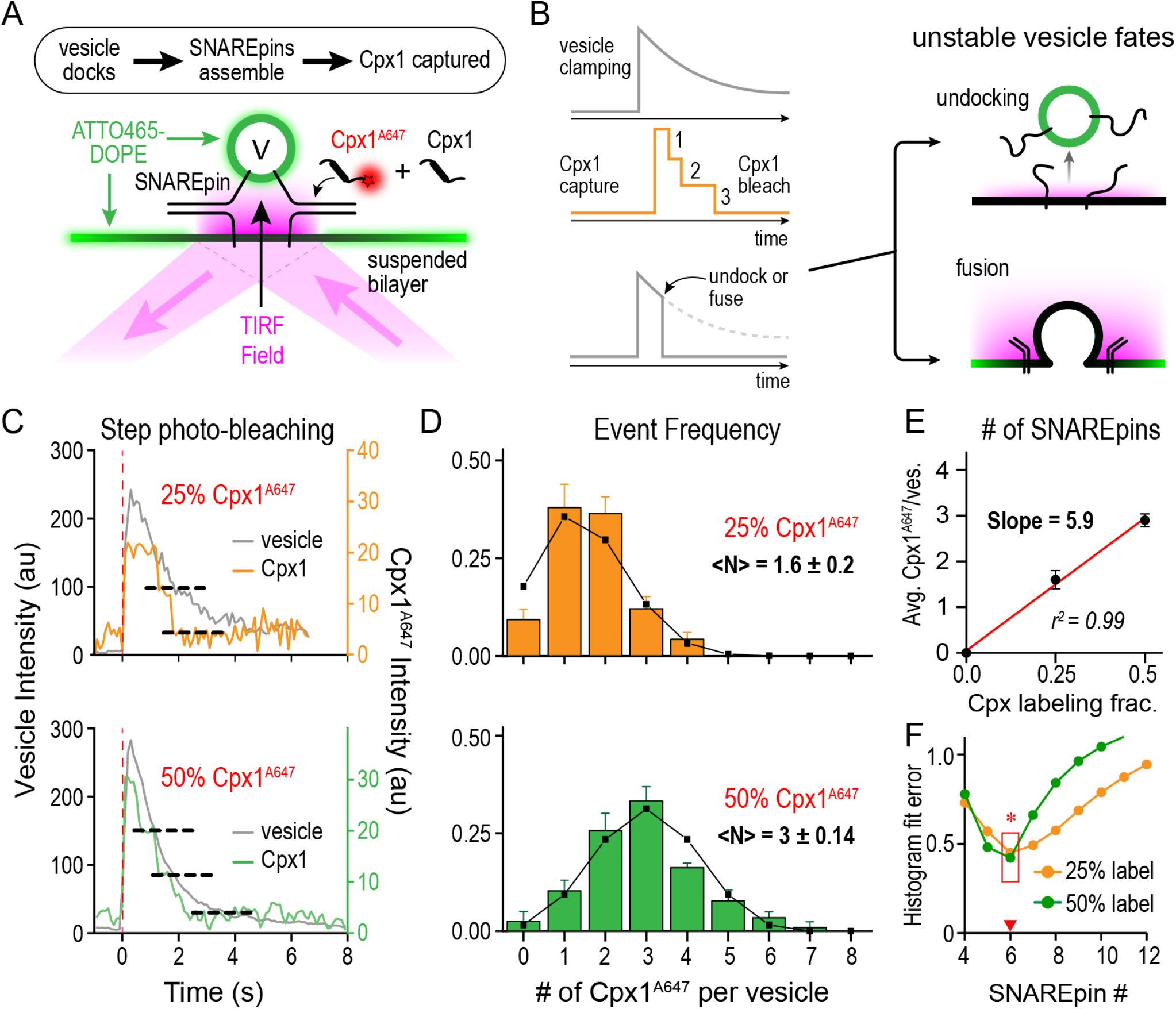
Precisely six SNAREpins are assembled under a clamped vesicle. **A.** Schematic for TIRF microscopy with single-molecule imaging. The same protein composition was used as in Fig. 3 except that both the suspended bilayer and vesicle were labeled with ATTO465-DOPE and Cpx1 labeled with Alexa Fluor 647 (Cpx1^A647^) was mixed with unlabeled Cpx1 at a ratio of either 1:3 (25% labeling) or 1:1 (50% labeling) for a total concentration of 2 µM. **B.** Two-color TIRF imaging was used to monitor vesicle arrival in the TIRF field as a rapid increase in fluorescence followed by a slow exponential decay due to bleaching (*top gray curve*) of immobile (clamped) vesicles. A small number of labeled Cpx1^A647^ molecules were monitored in the red channel while continuous bleaching produced step decays in Cpx1^A647^ fluorescence (*orange curve*) that were used to count Cpx1^A647^ molecules immobilized on the clamped vesicle. In some cases, immobile vesicle fluorescence would disappear within a single imaging frame (< 100 msec), indicating that the vesicle either fused or undocked and left the TIRF field. **C.** (*top*) Representative trace for a clamped vesicle (*gray*) that captured two Cpx1^A647^ molecules (*orange*) using 25% labeled Cpx1. (*bottom*) Representative trace for a clamped vesicle (*gray*) that captured three Cpx1^A647^ molecules (*green*) using 50% labeled Cpx1. **D.** Histograms of Cpx1^A647^ counts were plotted for 161 docked vesicles across three experiments using 25% Cpx1^A647^ (*orange*) and for 117 docked vesicles across three experiments using 50% Cpx1^A647^ (*green*), generating an average of 1.6 and 3.0 Cpx1^A647^, respectively. Predicted histograms using N = 6 binding sites for Cpx1^A647^ were generated using a binomial distribution with *p* = 0.25 (*black curve, top*) or *p* = 0.5 (*black curve, bottom*). **E.** The average number of Cpx molecules associated with a clamped vesicle plotted versus the fraction of labeled Cpx1 was fit to a line with a slope of 5.9, indicating that 6 Cpx1 molecules stably associate with a clamped vesicle. **F.** The 25% (*orange*) and 50% (*green*) histograms were fitted to a series of binomial models varying *N*, and the total fit error was plotted against *N* increasing from 4 to 12 (see **Methods** for details). The best fits in both cases occurred with *N* = 6. Data are presented as mean ± SEM.

The sole natural cysteine in Cpx1 (C105) was tagged with Alexa 647 (Cpx1^A647^) and mixed with unlabeled Cpx1 at a ratio of either 1:3 (25% label) or 1:1 (50% label) while keeping the total Cpx1 concentration at 2 μM to match the single-vesicle docking/fusion conditions described in **Fig. 3** while also keeping labeled Cpx1 within an optimal range for accurate single-molecule detection and counting. Importantly, the labeled cysteine does not participate in SNARE binding and resides more than 30 residues C-terminal to the Cpx1 central helix. Munc18-1/Syntaxin 1A complex and palmitoylated SNAP25 were reconstituted into GUVs, which were burst on a silicon surface to form a suspended bilayer as described earlier. Vesicles containing VAMP2 and Syt1 were fluorescently labeled using 1% DOPE-atto465 lipids and added to the supported bilayer together with the Cpx1/Cpx1^A647^ mixture, DHG, and wild-type Munc13C. In these TIRF experiments, the suspended bilayer also contained ATTO465-DOPE lipids to enable proper localization of the TIRF field, but during image acquisition, these fluorophores were steadily bleached to generate a low fluorescence background required for detection of vesicle arrival.

Tightly docked vesicles (hereon referred to as ‘clamped’ vesicles) were detected by a rapid increase in vesicle fluorescence intensity as the vesicle entered the TIRF field and became immobilized on the suspended bilayer followed by a slow exponential decay due to photobleaching of vesicular ATTO465 fluorophores (**Fig. 4C**). For wild-type Munc13, the majority of vesicles remained clamped during the entire image acquisition time window before bleaching finally eliminated the vesicle fluorophores (dwell time > 10 sec). Based on the confocal experiments with docked vesicles described above, the dwell time of clamped vesicles was likely much longer than 10 seconds, but photobleaching in the TIRF field prevented an accurate determination of the true dwell time. Under these conditions, Cpx1^A647^ arrived either simultaneously with or rapidly after vesicle clamping (<100 msec, limited by image acquisition rate). The number of captured Cpx molecules for each clamped vesicle was determined by analyzing the step-wise photobleaching time course (**Fig. 4C**). With 25% labeling, typically one or two Cpx1^A647^ molecules were captured upon vesicle arrival, whereas with 50% labeling, three Cpx1^A647^ molecules were captured (**Fig. 4C-D**). As only a fraction of Cpx1 molecules were fluorescently tagged, the Cpx1^A647^ signal generally corresponds to a larger number of total bound Cpx1 molecules. For individual trials however, binding is a stochastic process and the precise number of total bound Cpx1 molecules cannot be specified based solely on Cpx1^A647^ fluorescence. For example, in the presence of near saturating total Cpx1, given a 25% chance of Cpx1^A647^ capture per Cpx-bound SNAREpin and the relatively small number of total assembled SNAREpins per vesicle, one would expect 10-20% of clamped vesicles to be completely unlabeled by Cpx1^A647^ according to binomial statistics (eg. Prob(0 labels) = *(1 - f)^N^* where *f* is the fraction of labeled Cpx1 and N is the number of assembled SNAREpins assuming a range of *N = 5 – 8*). By contrast, with 50% labeling, the number of unlabeled clamped vesicles would be expected to fall to less than 2%. Indeed, we observed ∼10% and 2.5% unlabeled clamped vesicles for 25% and 50% Cpx1^A647^ labeling, respectively (**Fig. 4D**). Critically, both the 25% and 50% distributions of Cpx1 counts were well-described by binomial distributions with N = 6 ± 1 stable SNAREpins available to capture Cpx1 (**Fig. 4E-F**). Thus, in TIRF mode using wild-type Munc13C, individual clamped vesicles rapidly captured of Cpx1 onto six SNAREpins with 100 msec of vesicle docking, potentially enforced by the sixfold symmetry of the Munc13C assemblies.

### Vesicle docking and the assembly of six SNAREpins require Munc13C oligomerization

To test the hypotheses that vesicle clamping and the subsequent assembly of six SNAREpins were catalyzed by oligomerized Munc13C, we employed single-molecule TIRF Cpx1 counting using trimer and lateral Munc13 variants. In striking contrast to wild-type Munc13C, many of the vesicles docked by the trimer and lateral variants rapidly undocked or fused within seconds after immobilizing (**Fig. 5A-B**). Specifically, more than 65% of the vesicles docked by the trimer variants were unstable with vesicle fluorescence abruptly disappearing within 2-3 seconds of clamping, and the lateral FI/NN variant also exhibited a large increase in instability although less than the trimer variants (**Fig. 5A-B**). Given the threefold enhancement of spontaneous fusion observed by confocal imaging using the 2xR and 2xD trimer variants, many of these unstable events in TIRF imaging likely corresponded to spontaneous fusion. However, the high undocking rate observed in the lateral FI/NN variant by confocal imaging suggests that undocking accounted for most of the unstable events observed with lateral mutants. Another prominent difference observed was a pronounced slowing of the vesicle capture rate with both the trimer and lateral mutants requiring >15-fold longer time interval to clamp vesicles after they entered the TIRF field. Specifically, using wild-type Munc13C, vesicles entered the clamped state in less than 100 msec of nearing the suspended bilayer whereas clamping required ∼ 1.5 seconds using the trimer and lateral variants (**Fig. 5C-D**). And for events classified as unstable, the two trimer variants and the lateral FI/NN variant all exhibited similar vesicle dwell times of ∼ 2 seconds following entry into the clamped state. Thus, the clamped state catalyzed by Munc13 oligomer mutants was fundamentally volatile and vesicles progressed either to spontaneous fusion or undocked depending on which oligomer was destabilized. Unlike wild-type Munc13C where six stable SNAREpins were reliably and rapidly formed, vesicles docked by the trimer 2xR and 2xD variants as well as the lateral FI/NN and lateral quad variants formed only one or two stable SNAREpins based on 65-75% of observed docked vesicles failing to capture Cpx1 using 25% labeled Cpx1^A647^ (**Fig. 5F-I**). With destabilized trimers or lateral hexamers, the lower probability of forming and maintaining a functional oligomer prevented the stable assembly of 3 or more SNAREpins. In sum, irrespective of their differences, both classes of Munc13C mutations prolonged the time required to clamp vesicles, decreased the clamped-state dwell-time, and largely abolished stable SNAREpin formation, indicating requirements for both the trimer and lateral interfaces in these processes.

**Figure 5.**
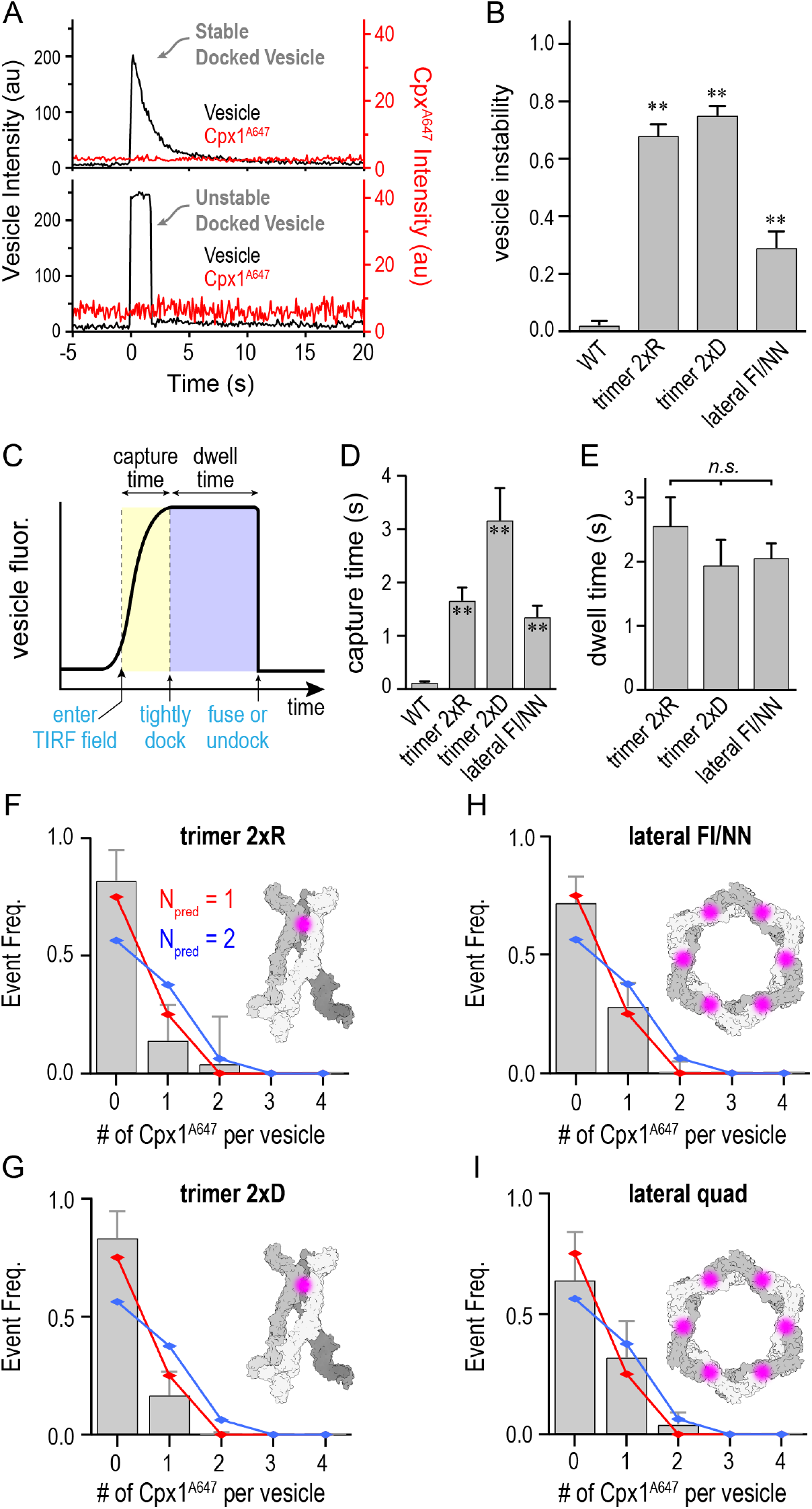
Vesicle docking and SNAREpin assembly are impaired by Munc13C trimer and lateral interface mutations. **A.** Example trace for a stable docked vesicle (*top, black*) and an unstable docked vesicle (bottom, black). In both cases, no Cpx1^A647^ molecules were detected (*red*). **B.** Vesicle instability was defined by the fraction of total vesicle docking events classified as unstable and plotted for experiments using wild-type Munc13C (WT), two trimer variants (2xR and 2xD), and one lateral hexamer variant (FI/NN). All interface mutations destabilized vesicle docking by a factor of 15 to 35-fold. **C.** For unstable docking events, the docking time course was partitioned into a capture phase as the vesicle entered the TIRF field but remained mobile (capture time) and a clamped phase where the vesicle was immobilized (dwell time). **D.** The mean capture time is plotted for wild-type Munc13C (WT) as well as three interface mutants (trimer 2xR, trimer 2xD, and lateral FI/NN). WT Munc13C captures vesicles 15 to 30-fold faster than the interface mutants. **E.** The mean dwell time is plotted for the same three interface mutants. Note that WT dwell time is longer than the duration of the experiment. Histograms for the number of Cpx molecules immobilized on clamped vesicles are shown for trimer 2xR (n=136 docked vesicles) (**F**), trimer 2xD (n=109) (**G**), lateral FI/NN (n=111) (**H**), and lateral quad (n = 177) (**I**). Binomial distributions with N = 1 (*red*) and N = 2 (*blue*) are superimposed on all four histograms for comparison. Data are presented as mean ± SEM. Data were compared using one-way ANOVA and the Tukey-Kramer analysis for multiple comparisons. ** *p < 0.01*.

### Disrupting the trimer and lateral interfaces impairs nervous system function and neurotransmitter secretion

Impairment of SNAREpin assembly, vesicle docking, and calcium-triggered vesicle fusion in the trimer and lateral interface variants of Munc13C support an important role for the oligomeric assemblies of Munc13C observed in CryoET. To further explore the functional impact of destabilized Munc13 oligomers in vivo, we examined the trimer and lateral interface variants using *C. elegans* as a model nervous system. Nematodes lacking the Munc13-1 ortholog UNC-13 are immobile with nearly complete loss of chemical synaptic transmission, and expression of the long UNC-13 isoform (UNC-13L) under a neuronal promoter has been shown to restore synaptic function ^19–23^. To disrupt the homologous trimer interface in worm UNC-13, we generated rescuing trimer 2xR and trimer 2xD UNC-13L constructs harboring D1459R/E1470R and K1585D/K1590D mutations, respectively (**Fig. 1C**). For the lateral interface, where only the hydrophobic component was conserved in *C. elegans*, the lateral FI/NN UNC-13L rescue construct contained I1283N/L1322N mutations (hereon referred to as lateral IL/NN) (**Fig. 2C**). Null mutant animals expressing the trimer 2xR or lateral IL/NN variants of UNC-13L exhibited more spontaneous movement than the *unc-13* null mutant. However, both mutant variants significantly impaired compared to null mutants expressing wild-type UNC-13L with 64% and 48% reductions in average worm speed for the trimer 2xR and lateral IL/NN variant, respectively (**Fig. 6A&B**). Thus, disruption of either oligomer interface in the worm ortholog significantly impaired locomotor behavior.

**Figure 6.**
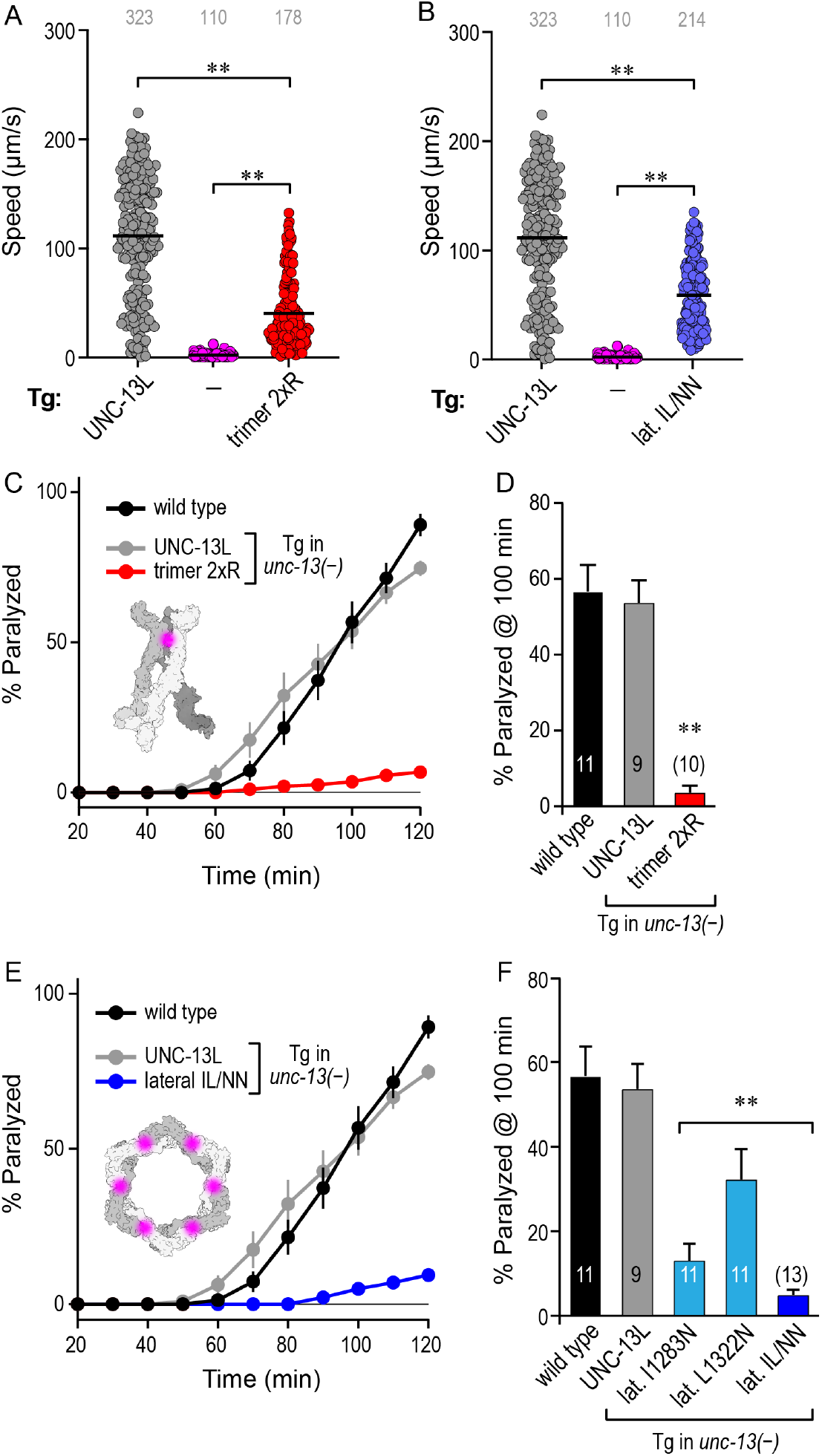
Perturbation of both the trimer and lateral structures impair nervous system function and neurotransmitter secretion. Spontaneous locomotion speed for *unc-13(s69)* null mutants (–, *magenta*), null mutants expressing a full-length UNC-13L transgene (UNC-13L, *gray*), null mutants expressing a trimer variant (trimer 2xR, *red*) (**A**), and null mutants expressing a trimer variant (trimer 2xR, *blue*) (**B**). Paralysis time course (**C**) and average percent paralyzed at 100 min (**D**) on 1 mM aldicarb for wild type (*black*) and *unc-13* null mutants expressing either wild-type UNC-13L (*gray*) or the UNC-13L trimer 2xR variant (*red*). **E.** Paralysis time on 1 mM aldicarb for wild type (*black*) and *unc-13* null mutants expressing either wild-type UNC-13L (*gray*) or the UNC-13L lateral IL/NN variant (*blue*). **F.** Average percent paralyzed at 100 min for wild type (*black*), null mutants expressing UNC-13L (*gray*), null mutants expressing a single lateral interface mutation in either MUN H7 (I1283N, *light blue*) or H8 (L1322N, *light blue*), and null mutants expressing the lateral IL/NN variant (*dark blue*). Data are mean ± SEM. For statistical comparisons. ** *p < 0.01* compared to wild type. Data were compared using one-way ANOVA and the Tukey-Kramer analysis for multiple comparisons.

As a more specific assay of neuromuscular synaptic function, sensitivity to the cholinesterase inhibitor, aldicarb, was monitored in the transgenic animals. Acetylcholine (ACh) is rapidly cleared from the neuromuscular junction (NMJ) by synaptic acetylcholinesterase enzymes (AChE), and pharmacological inhibition of AChE causes a buildup of ACh, leading to paralysis with a characteristic time course. Previous work has demonstrated that defects in synaptic transmission cause slower paralysis kinetics and resistance to aldicarb ^22,24–28^. Consistent with in vitro measures of Munc13C function in the trimer 2xR variant, trimer 2xR transgenic animals were markedly resistant to aldicarb, exhibiting little to no paralysis after 100 minutes of aldicarb exposure (**Fig. 6C&D**). Finally, the lateral IL/NN transgenic animals also displayed profound resistance to aldicarb treatment, similar to the trimer 2xR animals (**Fig. 6E&F**). Notably, even a single point mutation in helix H7 (I1283N) was sufficient to produce a significant impairment in ACh secretion by this assay. Overall, the spontaneous locomotion and aldicarb sensitivity assays indicated that disruptions in either the trimer or lateral interface were sufficient to impair neurotransmitter secretion at the *C. elegans* NMJ and bolster the conclusions drawn from our in vitro assays of homo-oligomeric Munc13C function.

## Discussion

We previously identified two Munc13C oligomers and proposed that these distinct assemblies reflect a key feature of the topological organization of the release apparatus. Here, we designed several Munc13C variants to disrupt either the trimer or lateral hexamer as monitored by CryoET through point mutations in the MUN domain while leaving overall protein stability intact. In vitro, perturbations in both oligomer types markedly slowed entry into the clamped-vesicle state and impaired calcium-triggered fusion of docked vesicles. In addition, the faithful assembly of six stable SNAREpins on clamped vesicles observed with wild-type Munc13C was abolished in the oligomer mutants. In vivo, both sets of oligomer mutations in the worm ortholog UNC-13 disrupted normal locomotor behavior and decreased ACh secretion. Notably, several differences between the trimer and lateral hexamer variants were also observed. First, a distinct Munc13C oligomer with a 2-fold symmetry emerged in the lateral FI/NN variant. This novel oligomer requires further investigation. Second, vesicles docked by this lateral variant rapidly undocked from membranes. Third, the destabilized trimer variant promoted spontaneous vesicle fusion while failing to support stable SNAREpin assembly. In sum, these findings reinforce the viewpoint that a precisely arranged assembly of Munc13 molecules coordinates discrete events including vesicle capture, tight docking, SNAREpin counting, and SNAREpin assembly and that each of these steps is required for efficient calcium-triggered fusion.

### Evidence for Munc13 multimerization

The results presented here bolster the hypothesis that Munc13 functions as a multimeric complex at release sites to promote SV docking and priming. Several in vitro studies of Munc13 have provided evidence for Munc13 oligomerization. Munc13-1 can homodimerize through its N-terminal C2A domain ^29^, but this region of the full-length protein was not included in the current study, indicating that even lacking this dimerization domain, the core domains of Munc13-1 can oligomerize. We have previously shown that Munc13C spontaneously clusters on membranes in vitro ^7^. Moreover, Munc13C can capture vesicles on supported bilayers with much higher efficiency when clusters of six or more monomers form, consistent with a cooperative process as well as with the six-fold symmetry of the Munc13C oligomers explored here ^6,7^. At synapses, several studies have described clustering of presynaptic Munc13-1 within the active zone. In particular, super-resolution imaging of Munc13-1 in hippocampal cultures revealed nanoassemblies of Munc13-1 containing an estimated 10-12 labeled monomers corresponding to a single release site ^30^. Freeze-fracture replica labeling in mouse cerebellar cortex and hippocampus also revealed similar small clusters of Munc13-1 associated with release sites ^31,32^. Interestingly, super-resolution imaging of the fly Munc13-1 ortholog Unc-13 in larval NMJ active zones detected prominent clustering with some evidence for subclusters containing three Unc-13 monomers, perhaps reflecting the homotrimeric state investigated here ^33^. Despite the daunting technical challenges of counting individual proteins on the tens of nanometers length scale within synaptic boutons, these studies collectively strengthen the viewpoint that multiple Munc13 monomers coalesce at single release sites to form functional units.

### Functional implications of Munc13 oligomers at release sites

What are the repercussions of Munc13 functioning as a multi-subunit protein complex rather than a monomer? Vesicle tethering and catalysis of SNARE assembly are two major roles previously proposed for Munc13, and both tasks are likely to profit from cooperativity and precision within a multi-subunit assembly ^4,5,34,35^. Membrane-bound Munc13C clusters in vitro capture and retain pure phospholipid vesicles only when six or more monomers are present and this steplike increase in avidity depends on the membrane-binding C-terminal C2C domain likely due to lashing several low affinity sites together into an effectively high affinity surface ^7^. Perhaps more critical than enhancing membrane binding, the ability to supervise a precise number of assembled SNAREpins may be the true payoff of Munc13 oligomers.

Work from many labs over the past two decades has demonstrated that only a small number of SNAREs are required to achieve membrane fusion and subsequent fusion pore expansion and yet the plasma membrane and vesicle membrane are inundated with a high density of SNARE proteins ^36–43^. Specifically, membrane fusion can be initiated by a single SNAREpin while at least three SNAREpins are thought to be required for successful pore expansion ^41–43^. Unique performance demands placed on the presynaptic fusion apparatus may have fashioned a relatively sophisticated molecular machine that can count and template a precise number of SNARE copies and integrate chemical information from multiple presynaptic proteins, small molecules, lipids, and calcium ions all while positioning these components at fixed locations between the plasma and vesicle membranes.

Given the likely scenario that any one SNARE assembly event may fail to produce an effective SNAREpin due to the stochastic behavior of single proteins, there is an obvious benefit of simultaneously templating six SNAREs within a single complex. For instance, binomial statistics reveal that to achieve a 95% chance of successful assembly of 3 SNAREpins when only 3 SNAREs are independently templated, the individual SNAREpins must assemble successfully with a probability greater than 98%. By contrast, six independent sites for SNAREpin assembly drops this required probability down to 70%. Furthermore, with modest positive cooperativity between assembling SNAREs due to their mechanical coupling within the Munc13 oligomer, the tolerance further drops to around 50% for individual SNAREpins.

An additional advantage of sequestering a small number of SNARE proteins into a highly confined and organized arrangement is a dramatic acceleration of SNARE assembly speed compared to an unorganized diffusion-limited alternative ^44^. The TIRF SNAREpin counting experiments presented here indicate that six SNAREpins are assembled in less than 100 msec upon vesicle docking, consistent with the observed rapid replenishment of synaptic release sites during activity that have an estimated time interval of 250 msec at the calyx of Held ^45^. And finally, the synchronicity of cooperative SNAREpin assembly can supply a greater force to bring membranes into close proximity and drive the fusion apparatus into a primed state. We propose that a succession of Munc13 oligomers guides a fixed number of SNAREs into assembled SNAREpins to achieve vesicle priming with both high speed and high fidelity. This molecular precision ensures a reliable and stereotyped fusion process within a synapse faced with a potentially broad range of activity levels.

### What is the sequence of events leading to SNARE assembly?

Mutations that disrupt either of the two distinct Munc13C assemblies were found to impair calcium-triggered fusion in vitro and neurotransmitter secretion in vivo. Thus, both oligomers reflect functionally important states of the pre-primed fusion machine and may also indicate specific steps in a stereotyped progression of Munc13 rearrangements that coordinate membrane positioning with SNARE templating. Although the experiments presented here do not specify a unique order to these steps, vesicle binding to the upright trimer followed by a transition to the lateral hexamer and ending with a flattening of the hexamer represents the most parsimonious ordering based on synaptic ultrastructural evidence for SVs moving from loosely tethered (20 nm gap between SV and plasma membrane) to tightly docked locations at the active zone (<10 nm gap) and the critical role played by Munc13 in these studies ^46,47^.

Numerous detailed sequential models of vesicle docking and SNARE assembly have been developed from biochemical, genetic, and physiological data and they are generally compatible an oligomeric framework for Munc13 function ^4,10,34,47–50^. One important feature of any sequential model is its treatment of directionality and reversibility as vesicles are moved through a series of states into a relatively stable primed state. SNARE zippering is likely to supply much of the free energy required to stabilize the primed state in all of these models, but SNARE proteins may promiscuously assemble in an unregulated fashion to trigger fusion while bypassing this state, a process partially antagonized by NSF/*α*SNAP-catalyzed SNARE disassembly ^51,52^. We noticed that the lateral hexagon interface occludes regions on the MUN domain indicated in Syntaxin 1 interactions proposed to expose its H3 SNARE domain and initiate SNARE assembly (**Fig. 2B**) ^53^. Additionally, the trimer 2xR mutations alter an aspartate proposed to interact with VAMP2 ^54^. Perhaps the interface-destabilizing mutations we characterize here also disrupt Syntaxin 1 opening and VAMP2 templating and thus also contribute to the poor performance of these Munc13 variants in vitro and in vivo. In this scenario, Munc13 oligomers could exert further control over SNARE assembly by coordinately hiding or exposing catalytic regions of the MUN domain. And SNARE binding after transitioning out of an inhibited state would antagonize the reverse reaction since the interface residues would be occupied by SNARE interactions. For instance, the enhanced undocking observed in the lateral FI/NN variant and elevated spontaneous fusion observed in the trimer variants described here may reflect different forms of decoupling of SNARE binding and vesicle docking. As the molecular details of SV docking, priming, and fusion continue to be elucidated, we anticipate that sequential and cooperative interactions between SNARE proteins and assemblies of core fusion proteins such as Munc13 will play a key part in accounting for the remarkable performance of the synaptic fusion apparatus.

### Potential roles for Munc13 oligomerization in synaptic plasticity and diversity

While the fundamental molecular mechanisms underlying synaptic transmission are largely identical across synapses within a nervous system as well as between the nervous systems of most animals, detailed studies of synaptic properties and their use-dependence have revealed that features such as synaptic strength and plasticity vary broadly across the mammalian brain ^32,55–58^. Numerous studies have implicated Munc13 isoforms as contributing to synaptic diversity and short-term presynaptic plasticities such as post-tetanic potentiation, augmentation, and use-dependent SV pool replenishment ^21,22,45,59–63^. Studies exploring presynaptic LTP in hippocampal mossy fibers and homeostatic plasticity in the fly NMJ reported changes in Munc13 distribution and abundance following plasticity induction ^33,64^. Karlocai and colleagues correlated Munc13-1 abundance and active zone distribution with synaptic properties across a population of hippocampal synapses revealing a broad heterogeneity of Munc13-1 abundance even within active zones possessing the same number of release sites ^32^. And Neher and colleagues hypothesized that shifts in a dynamic equilibrium between loosely docked and tightly docked states of SVs could account for some aspects of synaptic diversity and use-dependent plasticity ^58^. The central role of Munc13 in SV docking and its sensitivity to Ca^2+^ and lipids such as PI(4,5)P_2_ and DAG support the notion that Munc13 could play a part in multiple forms of presynaptic plasticity ^5,58,65^. And if Munc13 functions in larger homo-oligomeric complexes as proposed here, changes in monomer abundance as well as lipid and Ca^2+^ effects on oligomerization dynamics would be expected to impact synaptic strength via changes in SV docking, priming, and fusion. Notably, a human point mutation in the hinge region between the C2B and MUN domains of the human ortholog UNC13A was found to have a dominant gain-of-function synaptic phenotype perhaps resulting from destabilizing an autoinhibited state ^22,66^. Destabilizing the hinge region in a few monomers within a larger oligomeric UNC13A may be sufficient to release the autoinhibition and boost neurotransmitter release, possibly reflecting a mechanism normally used by Munc13 for use-dependent enhancement of synaptic strength.

### Conclusion and outlook

In this study, we tested the functional importance of two Munc13C oligomers previously identified by cryoEM tomography. While it is not yet established whether these particular molecular topologies occur at synaptic release sites, our results indicate that multimerization of Munc13 could have a significant functional impact on synaptic performance. In the context of other interacting active-zone and SV proteins in the highly crowded environment of a synapse, some aspects of these oligomeric structures and their associated SNARE stoichiometries may differ, but we suggest that Munc13 assemblies would still provide a means for counting SNAREs, driving rapid high-fidelity SNAREpin assembly, and efficiently organizing the associated priming machinery. Future studies will elucidate the structure of the fusion apparatus at synaptic release sites with increasing spatial resolution while more comprehensive in vitro vesicle fusion studies containing realistic SV and AZ compositions will continue to explore and refine these hypotheses.

## Materials and Methods

### Protein Constructs

The following cDNA constructs previously described ^13,67^ were used in this study: full-length VAMP2 (mouse His^6^-SUMO-VAMP2, residues 1-116); full-length SNAP25 (mouse His^6^-SNAP25b, residue 1-206); Synaptotagmin (rat Synaptotagmin1-His^6^, residues 57-421); Complexin (human His^6^-Complexin 1, residues 1-134) and Munc13 C1-C2B-MUN-C2C domain (rat His^12^-Munc13-1, residues 529-1735 with residues 1408–1452 replaced by the sequence EF and 1532-1550 deleted). We generated a new expression clone in the pET-duet vector to express and purify the Munc18-1/Syntaxin 1A complex (rat His^6^-SUMO Munc18/Syntaxin1). Phusion High Fidelity Mastermix (New England Biolabs, Ipswich, MA) was used to generate variants in Munc13 (trimer 2xR, trimer 2xD, trimer 3xD, and lateral FI/NN).

Lipids including 1, 2-Dioleoyl-sn-glycero-3-phosphocholine DOPC, 1, 2-Dioleoyl-sn-glycero-3-phospho-L-serine (DOPS), L-α-phosphatidylinositol-4, 5-bisphosphate (Brain PIP2) and 1, 2-Dioctadecanoyl-sn-glycerol (DAG) were purchased from Avanti Polar Lipids (Alabaster, AL). Fluorescent lipids ATTO465-DOPE and ATTO647N-DOPE were purchased from ATTO Tec (Siegen, Germany). 1, 2-Dihexanoyl-sn-glycerol was purchased from Cayman Chemicals (Ann Arbor, MI). TCEP HCl were purchased from Thermo Fisher (Waltham, MA).

### Protein Expression and Purification

All SNARE and associated proteins were expressed and purified as described previously ^15,68^. Briefly, proteins were expressed in Escherichia coli strain BL21(DE3) or Rosetta 2 (Novagen, Darmstadt, Germany) and cells were lysed with a cell disruptor (Avestin, Ottawa, Canada) in high salt buffer containing 400 mM KCl, 50 mm HEPES, 10% glycerol, pH 7.4, supplemented 2% Triton-X 100, 1 mm Tris (2-carboxyethyl) phosphine hydrochloride (TCEP), and 1 mm phenylmethylsulfonyl fluoride. Samples were clarified using a 45 Ti rotor (Beckman Coulter, Brea, CA, USA) for 30 minutes at 35, 000 rpm and incubated with Ni-NTA agarose (Thermo Fisher, Waltham, MA, USA) for 4–16 hours at 4 °C. The resin was subsequently washed (2 column volumes) in the wash buffer (high salt buffer supplemented with 1mM TCEP, 32 mM imidazole) with no detergent (Complexin and SNAP25) or 1% Octylglucoside (VAMP2 and Synaptotagmin) or 1% Triton X-100 (Munc18/Syntaxin). The proteins were either eluted with 300 mM Imidazole (Synaptotagmin) or cleaved of the resin with Thrombin (SNAP25 and Complexin) or SUMO protease (VAMP2 and Munc18/Syntaxin) in high salt buffer for 2 hours at room temperature. SNAP25 and Complexin proteins were further purified using gel filtration (Superdex75 Hi-load column, GE Healthcare, Chicago, IL), and Synaptotagmin-1 protein was subjected to the anion exchange column (MonoS, GE Healthcare, Chicago, IL) to remove nucleotide contaminants. The peak fractions were pooled and concentrated using filters of appropriate cutoffs (EMD Millipore, Burlington, MA, USA). SNAP25 was then palmitoylated using a 20-fold excess of Palmitoyl Coenzyme A in HEPES buffer supplemented with 1% TritonX-100 for 30 minutes at room temperature with gentle mixing.

Munc13C was purified as described previously ^6^. Briefly, Munc13C was expressed in ExpiHEK-293 cell cultures using ExpiFectamine as a transfection reagent (Thermo Fisher, Waltham, MA). Pellets were resuspended in lysis buffer and lysed using a Dounce homogenizer. The sample was clarified using at 35, 000 rpm for 30 minutes at 4 °C and the supernatant incubated overnight with Ni-NTA beads, in the presence of DNAse 1, RNAse A, and Benzonase to remove nucleotide contamination. The protein was further washed in the lysis buffer (without Triton-X 100) before being cleaved with PreScission protease for 2 hours at room temperature. The eluted proteins were further purified via gel filtration (Superdex 200, GE Healthcare Chicago, IL, USA). In all cases, the protein concentration was determined using a Bradford Assay (Bio-Rad, Hercules, CA, USA), with BSA as a standard, and protein purity was verified using SDS/PAGE analysis with Coomassie stain. All proteins were flash-frozen and stored at −80 °C for long-term storage.

### Suspended Bilayer and Vesicle Preparation

For the suspended bilayer, we approximated the pre-synaptic membrane physiological composition with 81% DOPC, 15% DOPS, 3% PIP2 and 1% ATTO465-PE for visualization. The lipids were mixed and dried under N_2_ gas and desiccated under vacuum. Bilayer samples were rehydrated with Munc18/Syntaxin1 and palmitoylated SNAP25 (1:1600 protein: lipid input ratio) in 5x buffer (125 mM HEPES, 600 mM KCl, 1 mM TCEP, pH 7.4) supplemented with 2% TritonX-100 for thirty minutes. Samples were then mixed directly with SM-2 Biobeads (Bio-Rad, Hercules, CA) for 30 minutes with gentle agitation. The samples were further dialyzed overnight in 5X buffer without detergent using a regenerated cellulose membrane (Spectrum Labs) on a flow dialysis with a 6-8 kDa molecular weight cut off.

Lipid bilayers were created by drying and rehydrating membranes to form GUVs as previously described ^13^. Briefly, 4 µL drops of Munc18-1/Syntaxin 1 + palymitoylated SNAP25 containing proteoliposomes were dried on a clean Mattek dish and rehydrated twice. In the second rehydration, the sample was diluted 5X to 20 µL with distilled water and then added to a cleaned silicon chip containing 1x buffer (25 mM HEPES, 120 mM KCl, 1 mM TCEP, pH 7.4) supplemented with Mg^2+^ (5 mM). After a 20-minute interval to allow bilayer formation, the bilayer was extensively washed with 1x buffer, and the fluidity of the lipid bilayer was verified using fluorescence recovery after photobleaching (FRAP) using the ATTO465 fluorescence.

For TIRF experiments, we created the suspended bilayer using a silicon chip, with buffer supplemented with 45% OptiPrep gradient media for index matching ^69^. Optiprep on the top was washed out with 1X buffer after bilayer formation. Bilayers contained 0.1% ATTO465 and were photobleached with 100% laser power after verifying they had formed. For small-unilamellar vesicle preparation, we approximated the synaptic vesicle lipid composition using 83% DOPC, 15% DOPS, 2% ATTO647N-PE. The samples were dried under N_2_ gas and desiccated under vacuum. Lipids were rehydrated with VAMP2 (1:100) and Synaptotagmin1 (1:250) in buffer (140 mM KCl, 50 mM HEPES, 1 mM TCEP, pH 7.4) supplemented with 1% Octyl β-glucoside. After 30 minutes of mixing, samples were rapidly diluted 3X below CMC and allowed to sit for another 30 minutes before being dialyzed overnight buffer without detergent. The samples were subjected to additional purification on the discontinuous Nycodenz gradient.

### Confocal single-vesicle fusion assays

Single vesicle assays were performed as described previously with a few modifications ^13^. DHG was added to the pre-formed bilayer and incubated for 5 minutes. Vesicles (50 µM lipids) were added from the top using a pipette and allowed to interact with the bilayer for 3 minutes. We used ATTO647-labeled DOPE fluorescence to track the fate of the individual vesicles. All vesicles that attach to the suspended bilayer during the 3-minute observation period were scored as docked. Initially, all docked vesicles exhibited diffusive mobility and subsequently transitioned to an immobile tightly-docked state. Among these mobile and immobile vesicles, some underwent spontaneous fusion, evident by a burst of fluorescence intensity followed by a rapid decrease. Furthermore, we observed instances of undocking, where vesicle fluorescence disappeared without a fusion burst, primarily during the mobile phase. Undocking events were infrequent once vesicles reached the immobile state. After the initial 3 min interaction phase, the excess vesicles in the chamber were removed by buffer exchange (3X buffer wash) and 100 mM CaCl_2_ was added from the top to monitor the effect of Ca^2+^ on the docked vesicles.

### TIRF single-vesicle Cpx-counting assays

All the complexin counting experiments underneath docked vesicles were performed with TIRF microscopy (Nikon) on a reconstituted planar suspended bilayer on the silicon surface. The method to create a suspended bilayer compatible with single-molecule imaging was described in detail elsewhere ^69^. Briefly, a suspended membrane (using the composition described above) was prepared using the osmotic shock protocol and burst onto freshly plasma-cleaned Si/SiO_2_ chips containing 5 µm diameter holes in presence of HEPES buffer (25 mM HEPES, 140 mM KCl, 1 mM DTT) supplemented with index matching 45% Optiprep(TM) (Stemcell Technologies) and 5 mM MgCl_2_ ^70^. After waiting 20 minutes, the suspended bilayer was extensively washed with HEPES buffer containing 1 mM MgCl_2_. Before adding any vesicles, the bilayer was completely bleached and then 2 µM total complexin (with corresponding labeled protein) was added and mixed well. Both the vesicle and labeled complexin were simultaneously monitored with 488 and 633 nm solid-state lasers respectively using dual viewer 2. All the data were analyzed with ImageJ. The number of labeled complexin molecules underneath a vesicle was calculated from the measured step-wise photobleaching. Cpx1^A647^ counting distributions were compared to binomial distributions: probability 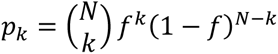 where *N* is the number of binding sites (SNAREpins) and *f* is the fraction of total Cpx1 labeled with a fluorophore. The total error between the measured distribution *d_k_* and a binomial distribution *p_k_* for *f = 0.25* or *f = 0.5* and a range of values for *N* was computed as 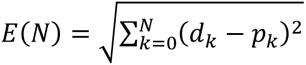 (**Fig. 4F**).

### C. elegans strains

Strains were maintained and genetically manipulated as previously described ^71^. Animals were raised at 20°C on nematode growth media seeded with OP50. Primary strains used: N2, *unc-13(s69)*. JSD1392: *tauEx574;unc-13(s69)* = P*_snb-1_*::UNC-13L wild-type rescue construct. JSD1385: *tauEx571;unc-13(s69)* = P*_snb-1_*::UNC-13L(D1459R;E1470R) 2xR trimer variant. JSD1413: *tauEx579;unc-13(s69)* = P*_snb-1_*::UNC-13L(L1322N). JSD1414: *tauEx580;unc-13(s69)* = P*_snb-1_*::UNC-13L(I1283N). JSD1415: *tauEx581;unc-13(s69)* = P*_snb-1_*::UNC-13L(I1283N;L1322N) lateral IL/NN variant.

### Aldicarb resistance assay

To measure aldicarb sensitivity in *C. elegans*, approximately 20 animals were placed on agar plates containing 1 mM aldicarb (CarboSynth) as described previously ^26,72^. Animals were scored for paralysis every 10 minutes for 2 hours for each genotype. The experimenter was blind to the genotype, and each genotype was assayed at least 10 times. Paralysis curves for each genotype were generated by averaging time courses.

### Spontaneous locomotion

For *C. elegans* locomotion, approximately 40 worms of each genotype were moved to agar plates with no bacterial food using M9 buffer one hour before tracking. One-minute videos of the spontaneous movement were captured using an ORCA-05G CCD camera (Hamamatsu) and an Olympus SZX16 stereomicroscope. The movement was tracked by a custom object tracking software on Matlab ^73^. Worms that moved 30 seconds or less were not included in the data set. Each genotype has an n between 100-300 individual worms that were tracked. Speeds for each genotype were pooled and analyzed by one-way ANOVA and post hoc analysis across all genotypes.

### Lipid Membrane Preparation for Cryo-electron Tomography

Vesicles were prepared with a lipid composition consisting of DOPC/DOPS/PIP2 in a molar ratio of 14/80/6. The lipid stocks were mixed in a chloroform with addition of 20 μL methanol to dissolve PIP2 and the solvent was evaporated under N2 gas followed by vacuum drying for 1 h. The resulting dried lipid film was rehydrated for 1 h at room temperature with constant vortexing in buffer containing 20 mM MOPS pH 7.4, 150 mM KCl, 1 mM EDTA, and 0.5 mM TCEP at a final lipid concentration of 1 mM. After rehydration, the mixture was sonicated for 5 min using a bath sonicator (Branson Ultrasonics). To remove large aggregates of lipid membranes, the prepared solution was stored overnight at 4 °C. Vesicles were separated from sedimented aggregates and used for crystallization experiments.

### Protein Crystallization

The protocol resulted in unilamellar vesicles with broad range in sizes. Crystals were formed immediately before the freezing by mixing 1 μM Munc13C with 100 μM lipid membranes in 1:1 (vol/vol) ratio, total volume of 20 μL. Once mixed, samples were incubated at room temperature for 5 min before freezing.

### Electron Microscopy Sample Preparation and Data Acquisition

Samples were vitrified using a Vitrobot Mark IV (Thermo Fisher Scientific) held at 8 °C with 100% humidity. Bovine serum albumin (BSA)-coated, 10-nm Gold Tracer beads (Aurion) were added immediately prior to vitrification to the samples as fiducial markers. Samples (2.5 μL) were applied to freshly glow-discharged 200 mesh Lacey Formvar/carbon grids directly in the blotting chamber, and grids were blotted for 5 s with blot force −1 and then plunge frozen in liquid ethane cooled by liquid nitrogen. Initially, samples were screened with a Glacios Cryo TEM 200 kV (Thermo Fisher Scientific) equipped with a K2 Summit direct electron detector (Gatan). Low- and medium-resolution montages were acquired for a complete grid and selected squares overview. The K2 camera was used in counting mode, and detector dark and gain references were collected prior to each data acquisition session. Cryoelectron tomograms were acquired using SerialEM with a bidirectional scheme from 0°; - 51° to +51°,3°increment, 35 frames, and 54 e^-^/ Å^2^ total dose. The nominal magnification was 13,500× resulting in a pixel size of 3.019 Å. Nominal defocus was set to −4 μm. Collected tilt series were selected for further processing based on the visual appearance of the crystal, resulting in 10 total tomograms for initial 3D reconstruction. The final dataset was collected using a 300 kV Titan Krios G2 transmission electron microscope (Thermo Fisher Scientific) equipped with GIF Quantum LS energy filter mounted in front of a K3 Summit direct electron detector (Gatan). The slit width of the filter was set to 20 eV. Filter tuning was done using Digital Micrograph software (Gatan). The tilt-series images were acquired using serial-EM at a nominal magnification of 42,000× (corresponding to a calibrated physical pixel size of 2.1 Å) using dose symmetric scheme (78) with tilt range ± 51° and 3° increment. The tilt images were acquired as 11,520 × 8,184 super-resolution movies of 12 frames for the first session or 14 frames for the second one per each tilt corresponding to a tilt series total dose ∼ 100 e^-^/ Å^2^ and ∼120 e^-^/ Å^2^, respectively. Defocus nominal range was set to −3.5 μm to −5 μm.

## Acknowledgements

We thank Dr. Shyam Krishnakumar for helpful discussions and members of the Rothman and Dittman labs for critically reading the manuscript. This work was supported by grants from NIH DK 027044 (JER) and NIH NS 116747 (JSD).

---

**Figure S1.**
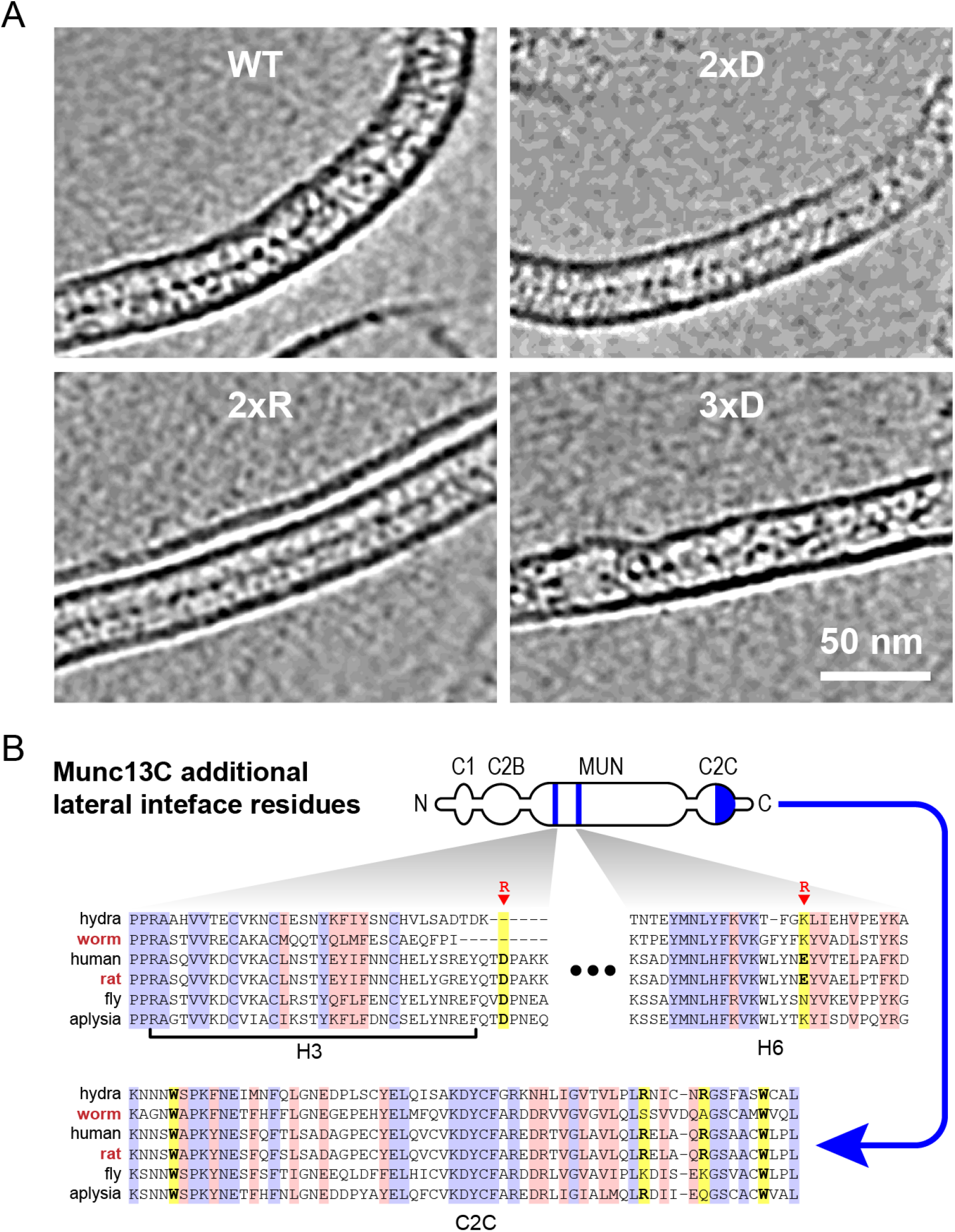
CryoET side views of trimer variants. **A.** Representative slices through the reconstructed cryo-electron tomograms (*side view*) to visualize Munc13C molecules residing between the two lipid bilayers for wild type (*top left*), trimer 2xD (top right), trimer 2xR (*bottom left*), and trimer 3xD (*bottom right*). In all cases, a thin protein density running between the two bilayers corresponds to Munc13C lateral hexamers. Scale bar is 50 nm. **B.** Sequence alignments for MUN H3 and H6 helices as well as the C2C domain with identical and similar residues highlighted in blue and red, respectively. D1039 (adjacent to H3) and D1139 (in H6) were mutated to R along with F1176N and I1215N in the lateral quad variant (*yellow*). The corresponding residues in C2C are also highlighted in yellow. Protein sequence accession numbers are given in Figure 1.

## References

1 Neher, E. & Sakaba, T. Multiple roles of calcium ions in the regulation of neurotransmitter release. Neuron 59, 861–872 (2008).

2 Sabatini, B. L. & Regehr, W. G. Timing of neurotransmission at fast synapses in the mammalian brain. Nature 384, 170–172 (1996). https://doi.org:10.1038/384170a0

3 Dittman, J. S. & Ryan, T. A. The control of release probability at nerve terminals. Nature Reviews Neuroscience 20 (2019).

4 Rizo, J. Molecular Mechanisms Underlying Neurotransmitter Release. Annu Rev Biophys 51, 377–408 (2022). https://doi.org:10.1146/annurev-biophys-111821-104732

5 Dittman, J. S. Unc13: a multifunctional synaptic marvel. Curr Opin Neurobiol 57, 17–25 (2019). https://doi.org:10.1016/j.conb.2018.12.011

6 Grushin, K., Kalyana Sundaram, R. V., Sindelar, C. V. & Rothman, J. E. Munc13 structural transitions and oligomers that may choreograph successive stages in vesicle priming for neurotransmitter release. Proc Natl Acad Sci U S A 119 (2022). https://doi.org:10.1073/pnas.2121259119

7 Li, F. et al. Vesicle capture by membrane-bound Munc13-1 requires self-assembly into discrete clusters. FEBS Lett 595, 2185–2196 (2021). https://doi.org:10.1002/1873-3468.14157

8 Xu, J. et al. Mechanistic insights into neurotransmitter release and presynaptic plasticity from the crystal structure of Munc13-1 C1C2BMUN. Elife 6 (2017). https://doi.org:10.7554/eLife.22567

9 Quade, B. et al. Membrane bridging by Munc13-1 is crucial for neurotransmitter release. Elife 8 (2019). https://doi.org:10.7554/eLife.42806

10 Camacho, M. et al. Control of neurotransmitter release by two distinct membrane-binding faces of the Munc13-1 C. Elife 10 (2021). https://doi.org:10.7554/eLife.72030

11 Shu, T., Jin, H., Rothman, J. E. & Zhang, Y. Munc13-1 MUN domain and Munc18-1 cooperatively chaperone SNARE assembly through a tetrameric complex. Proc Natl Acad Sci U S A 117, 1036–1041 (2020). https://doi.org:10.1073/pnas.1914361117

12 Liu, X. et al. Functional synergy between the Munc13 C-terminal C1 and C2 domains. Elife 5 (2016). https://doi.org:10.7554/eLife.13696

13 Sundaram, R. V. K. et al. Novel Roles for Diacylglycerol in Synaptic Vesicle Priming and Release Revealed by Complete Reconstitution of Core Protein Machinery. bioRxiv (2023). https://doi.org:10.1101/2023.06.05.543781

14 Ramakrishnan, S. et al. High-Throughput Monitoring of Single Vesicle Fusion Using Freestanding Membranes and Automated Analysis. Langmuir 34, 5849–5859 (2018). https://doi.org:10.1021/acs.langmuir.8b00116

15 Ramakrishnan, S., Bera, M., Coleman, J., Rothman, J. E. & Krishnakumar, S. S. Synergistic roles of Synaptotagmin-1 and complexin in calcium-regulated neuronal exocytosis. Elife 9 (2020). https://doi.org:10.7554/eLife.54506

16 Rothman, J. E., Grushin, K., Bera, M. & Pincet, F. Turbocharging Synaptic Transmission. FEBS Letters (2023).

17 Chen, X. et al. Three-dimensional structure of the complexin/SNARE complex. Neuron 33, 397–409 (2002).

18 Pabst, S. et al. Selective interaction of complexin with the neuronal SNARE complex. Determination of the binding regions. J Biol Chem 275, 19808–19818 (2000).

19 Richmond, J. E., Davis, W. S. & Jorgensen, E. M. UNC-13 is required for synaptic vesicle fusion in C. elegans. Nat Neurosci 2, 959–964 (1999).

20 Madison, J. M., Nurrish, S. & Kaplan, J. M. UNC-13 interaction with syntaxin is required for synaptic transmission. Curr Biol 15, 2236–2242 (2005).

21 Hu, Z., Vashlishan-Murray, A. B. & Kaplan, J. M. NLP-12 Engages Different UNC-13 Proteins to Potentiate Tonic and Evoked Release. J Neurosci 35, 1038–1042 (2015). https://doi.org:10.1523/jneurosci.2825-14.2015

22 Michelassi, F., Liu, H., Hu, Z. & Dittman, J. S. A C1-C2 Module in Munc13 Inhibits Calcium-Dependent Neurotransmitter Release. Neuron 95, 577–590.e575 (2017). https://doi.org:10.1016/j.neuron.2017.07.015

23 Liu, H. et al. The M domain in UNC-13 regulates the probability of neurotransmitter release. Cell Rep 34, 108828 (2021). https://doi.org:10.1016/j.celrep.2021.108828

24 Rand, J. B. & Russell, R. L. Molecular basis of drug-resistance mutations in C. elegans. Psychopharmacol Bull 21, 623–630 (1985).

25 Miller, K. G. et al. A genetic selection for Caenorhabditis elegans synaptic transmission mutants. Proc Natl Acad Sci U S A 93, 12593–12598 (1996).

26 Nurrish, S., Segalat, L. & Kaplan, J. M. Serotonin inhibition of synaptic transmission: Galpha(0) decreases the abundance of UNC-13 at release sites. Neuron 24, 231–242 (1999).

27 Martin, J. A., Hu, Z., Fenz, K. M., Fernandez, J. & Dittman, J. S. Complexin has opposite effects on two modes of synaptic vesicle fusion. Curr Biol 21, 97–105 (2011). https://doi.org:S0960-9822(10)01600

28 Wragg, R. T. et al. Evolutionary Divergence of the C-terminal Domain of Complexin Accounts for Functional Disparities between Vertebrate and Invertebrate Complexins. Front Mol Neurosci 10, 146 (2017). https://doi.org:10.3389/fnmol.2017.00146

29 Lu, J. et al. Structural basis for a Munc13-1 homodimer to Munc13-1/RIM heterodimer switch. PLoS Biol 4, e192 (2006). https://doi.org:10.1371/journal.pbio.0040192

30 Sakamoto, H. et al. Synaptic weight set by Munc13-1 supramolecular assemblies. Nat Neurosci 21, 41–49 (2018). https://doi.org:10.1038/s41593-017-0041-9

31 Rebola, N. et al. Distinct Nanoscale Calcium Channel and Synaptic Vesicle Topographies Contribute to the Diversity of Synaptic Function. Neuron 104, 693–710.e699 (2019). https://doi.org:10.1016/j.neuron.2019.08.014

32 Karlocai, M. R. et al. Variability in the Munc13-1 content of excitatory release sites. Elife 10 (2021). https://doi.org:10.7554/eLife.67468

33 Dannhäuser, S. et al. Endogenous tagging of Unc-13 reveals nanoscale reorganization at active zones during presynaptic homeostatic potentiation. Front Cell Neurosci 16, 1074304 (2022). https://doi.org:10.3389/fncel.2022.1074304

34 Rothman, J. E., Krishnakumar, S. S., Grushin, K. & Pincet, F. Hypothesis - buttressed rings assemble, clamp, and release SNAREpins for synaptic transmission. FEBS Lett 591, 3459–3480 (2017). https://doi.org:10.1002/1873-3468.12874

35 Padmanarayana, M. et al. A unique C2 domain at the C terminus of Munc13 promotes synaptic vesicle priming. Proc Natl Acad Sci U S A 118 (2021). https://doi.org:10.1073/pnas.2016276118

36 Hua, Y. & Scheller, R. H. Three SNARE complexes cooperate to mediate membrane fusion. Proc Natl Acad Sci U S A 98, 8065–8070 (2001). https://doi.org:10.1073/pnas.131214798

37 Han, X., Wang, C. T., Bai, J., Chapman, E. R. & Jackson, M. B. Transmembrane segments of syntaxin line the fusion pore of Ca2+-triggered exocytosis. Science 304, 289–292 (2004). https://doi.org:10.1126/science.1095801

38 Domanska, M. K., Kiessling, V., Stein, A., Fasshauer, D. & Tamm, L. K. Single vesicle millisecond fusion kinetics reveals number of SNARE complexes optimal for fast SNARE-mediated membrane fusion. J Biol Chem 284, 32158–32166 (2009). https://doi.org:10.1074/jbc.M109.047381

39 van den Bogaart, G., et al. One SNARE complex is sufficient for membrane fusion. Nat Struct Mol Biol 17, 358–364 (2010). https://doi.org:10.1038/nsmb.1748

40 Mohrmann, R., de Wit, H., Verhage, M., Neher, E. & Sorensen, J. B. Fast vesicle fusion in living cells requires at least three SNARE complexes. Science 330, 502–505 (2010).

41 Shi, L., et al. SNARE proteins: one to fuse and three to keep the nascent fusion pore open. Science 335, 1355–1359 (2012). https://doi.org:10.1126/science.1214984

42 Bao, H. et al. Dynamics and number of trans-SNARE complexes determine nascent fusion pore properties. Nature 554, 260–263 (2018). https://doi.org:10.1038/nature25481

43 Heo, P., Coleman, J., Fleury, J. B., Rothman, J. E. & Pincet, F. Nascent fusion pore opening monitored at single-SNAREpin resolution. Proc Natl Acad Sci U S A 118 (2021). https://doi.org:10.1073/pnas.2024922118

44 Mion, D., Bunel, L., Heo, P. & Pincet, F. The beginning and the end of SNARE-induced membrane fusion. FEBS Open Bio 12, 1958–1979 (2022). https://doi.org:10.1002/2211-5463.13447

45 Lipstein, N. et al. Munc13-1 is a Ca. Neuron 109, 3980–4000.e3987 (2021). https://doi.org:10.1016/j.neuron.2021.09.054

46 Imig, C. et al. The morphological and molecular nature of synaptic vesicle priming at presynaptic active zones. Neuron 84, 416–431 (2014). https://doi.org:10.1016/j.neuron.2014.10.009

47 Papantoniou, C. et al. Munc13- and SNAP25-dependent molecular bridges play a key role in synaptic vesicle priming. Sci Adv 9, eadf6222 (2023). https://doi.org:10.1126/sciadv.adf6222

48 Richmond, J. E., Weimer, R. M. & Jorgensen, E. M. An open form of syntaxin bypasses the requirement for UNC-13 in vesicle priming. Nature 412, 338–341 (2001).

49 Ma, C., Su, L., Seven, A. B., Xu, Y. & Rizo, J. Reconstitution of the vital functions of Munc18 and Munc13 in neurotransmitter release. Science 339, 421–425 (2013). https://doi.org:10.1126/science.1230473

50 Baker, R. W. et al. A direct role for the Sec1/Munc18-family protein Vps33 as a template for SNARE assembly. Science 349, 1111–1114 (2015). https://doi.org:10.1126/science.aac7906

51 Prinslow, E. A., Stepien, K. P., Pan, Y. Z., Xu, J. & Rizo, J. Multiple factors maintain assembled trans-SNARE complexes in the presence of NSF and αSNAP. Elife 8 (2019). https://doi.org:10.7554/eLife.38880

52 Brunger, A. T. et al. The pre-synaptic fusion machinery. Curr Opin Struct Biol 54, 179–188 (2019). https://doi.org:10.1016/j.sbi.2019.03.007

53 Yang, X. et al. Syntaxin opening by the MUN domain underlies the function of Munc13 in synaptic-vesicle priming. Nat Struct Mol Biol 22, 547–554 (2015). https://doi.org:10.1038/nsmb.3038

54 Wang, S. et al. Munc18 and Munc13 serve as a functional template to orchestrate neuronal SNARE complex assembly. Nat Commun 10, 69 (2019). https://doi.org:10.1038/s41467-018-08028-6

55 Dittman, J. S., Kreitzer, A. C. & Regehr, W. G. Interplay between facilitation, depression, and residual calcium at three presynaptic terminals. J Neurosci 20, 1374–1385 (2000).

56 O’Rourke, N. A., Weiler, N. C., Micheva, K. D. & Smith, S. J. Deep molecular diversity of mammalian synapses: why it matters and how to measure it. Nat Rev Neurosci 13, 365–379 (2012). https://doi.org:10.1038/nrn3170

57 Burkhardt, P. & Sprecher, S. G. Evolutionary origin of synapses and neurons - Bridging the gap. Bioessays 39 (2017). https://doi.org:10.1002/bies.201700024

58 Lin, K. H., Taschenberger, H. & Neher, E. A sequential two-step priming scheme reproduces diversity in synaptic strength and short-term plasticity. Proc Natl Acad Sci U S A 119, e2207987119 (2022). https://doi.org:10.1073/pnas.2207987119

59 Junge, H. J. et al. Calmodulin and Munc13 form a Ca2+ sensor/effector complex that controls short-term synaptic plasticity. Cell 118, 389–401 (2004). https://doi.org:10.1016/j.cell.2004.06.029

60 Lipstein, N. et al. Nonconserved Ca(2+)/calmodulin binding sites in Munc13s differentially control synaptic short-term plasticity. Mol Cell Biol 32, 4628–4641 (2012). https://doi.org:10.1128/mcb.00933-12

61 Lipstein, N. et al. Dynamic Control of Synaptic Vesicle Replenishment and Short-Term Plasticity by Ca2+-Calmodulin-Munc13-1 Signaling. Neuron 79, 82–96 (2013). https://doi.org:10.1016/j.neuron.2013.05.011

62 Lee, J. S., Ho, W. K., Neher, E. & Lee, S. H. Superpriming of synaptic vesicles after their recruitment to the readily releasable pool. Proc Natl Acad Sci U S A 110, 15079–15084 (2013). https://doi.org:10.1073/pnas.1314427110

63 Taschenberger, H., Woehler, A. & Neher, E. Superpriming of synaptic vesicles as a common basis for intersynapse variability and modulation of synaptic strength. Proc Natl Acad Sci U S A 113, E4548–4557 (2016). https://doi.org:10.1073/pnas.1606383113

64 Fukaya, R. et al. Increased vesicle fusion competence underlies long-term potentiation at hippocampal mossy fiber synapses. Sci Adv 9, eadd3616 (2023). https://doi.org:10.1126/sciadv.add3616

65 Silva, M., Tran, V. & Marty, A. Calcium-dependent docking of synaptic vesicles. Trends Neurosci 44, 579–592 (2021). https://doi.org:10.1016/j.tins.2021.04.003

66 Lipstein, N. et al. Synaptic UNC13A protein variant causes increased neurotransmission and dyskinetic movement disorder. J Clin Invest 127, 1005–1018 (2017). https://doi.org:10.1172/jci90259

67 Kalyana Sundaram, R. V., et al. Munc13 binds and recruits SNAP25 to chaperone SNARE complex assembly. FEBS Lett 595, 297–309 (2021). https://doi.org:10.1002/1873-3468.14006

68 Kalyana Sundaram, R. V., et al. Munc13 binds and recruits SNAP25 to chaperone SNARE complex assembly. FEBS Lett (2020). https://doi.org:10.1002/1873-3468.14006

69 Kalyana Sundaram, R. V., et al. Native Planar Asymmetric Suspended Membrane for Single-Molecule Investigations: Plasma Membrane on a Chip. Small 18, e2205567 (2022). https://doi.org:10.1002/smll.202205567

70 Motta, I. et al. Formation of Giant Unilamellar Proteo-Liposomes by Osmotic Shock. Langmuir 31, 7091–7099 (2015). https://doi.org:10.1021/acs.langmuir.5b01173

71 Brenner, S. The genetics of Caenorhabditis elegans. Genetics 77, 71–94 (1974).

72 Dittman, J. S. & Kaplan, J. M. Behavioral impact of neurotransmitter-activated G-protein-coupled receptors: muscarinic and GABAB receptors regulate Caenorhabditis elegans locomotion. J Neurosci 28, 7104–7112 (2008).

73 Ramot, D., Johnson, B. E., Berry, T. L., Jr., Carnell, L. & Goodman, M. B. The Parallel Worm Tracker: a platform for measuring average speed and drug-induced paralysis in nematodes. PLoS ONE 3, e2208 (2008). https://doi.org:10.1371/journal.pone.0002208

